# Co-option of Lysosomal Machinery for Sponge Biosilicification

**DOI:** 10.64898/2026.01.19.700231

**Authors:** Maitri Rangarajan-Paul, Scott A. Nichols, Martin Tresguerres

**Affiliations:** Scripps Institution of Oceanography, University of California San Diego, 9500 Gilman Drive, La Jolla, CA 92093, USA; University of Denver, Department of Biological Sciences, 2101 E. Wesley Ave SGM 203, Denver, CO 80210-5210

**Author notes:** Correspondence (M.R.P.), (M.T.).

## Abstract

Biomineralization evolved repeatedly across animals, resulting in novel strategies for buoyancy, locomotion, and defense. Sponges are the only metazoans that build a silica-based skeleton and the mechanisms underlying their biosilicification are relatively understudied compared to other major silicifiers. Here, we show that in freshwater sponges, cells specialized for biosilicification, termed sclerocytes, highly express lysosomal-associated genes, including the V-H^+^-ATPase (VHA) proton pump, oculocutaneous albinism type 2 (OCA2) anion channels, cathepsins, silicateins, TMEM55A/B, TMEM192, TMEM199, and other proteins involved in lysosomal maintenance and degradation. We demonstrate that VHA protein localizes to the sclerocyte silica deposition vesicle (SDV) and that VHA-dependent SDV acidification is essential for silica spicule formation. This function mirrors the role of VHA in diatom biosilicification and in biocalcification across a broad sweep of taxa. We also found that genes homologous to the plant silicon efflux transporter Lsi2 are highly upregulated in sclerocytes and confirmed their localization using Hybridization Chain Reaction Fluorescence In Situ Hybridization (HCR-FISH). These results corroborate phylogenetic and transcriptomic evidence that Lsi2 homologs are involved in sponge silicic acid transport. Data mining of previous studies revealed many of the sponge sclerocyte lysosomal-associated genes have been correlated with biomineralization across eukaryotes, including coccolithophores, bryozoans, crustaceans, mollusks, and mammals. As detoxification was a likely driver for the evolution of biomineralization, we propose that lysosomal machinery aided in detoxification by allowing sponges to sequester Si in acidified compartments to prevent its catastrophic intracellular precipitation. This lysosomal toolkit was likely independently co-opted for the formation of biomineralized structures in multiple species.

## Introduction

The emergence of biomineralization, the formation of mineral structures by living organisms, played a key role in the explosion of body plan diversity during the Cambrian (Smith and Harper 2013). Sponges are the earliest diverged branch of metazoans (Steenwyk and King 2025) and are the only metazoans that build a silica-based skeleton (Müller et al. 2009). The cellular and molecular mechanisms underlying their silicification process are much less understood than in diatoms and land plants, the other two major biosilicifiers. One of the proposed selective pressures for the evolution of skeleton formation is detoxification (Simkiss 1977; Simkiss 1989; Kempe and Kazmierczak 1994). Si(OH)_4_ and Ca^2+^ were present at high levels in the ocean during the pre-Cambrian/Cambrian boundary when the biomineralization began to emerge (Brennan et al. 2004; Marron et al. 2016). Given that their influx into cells and subsequent uncontrolled precipitation would severely disrupt cellular homeostasis (Orrenius et al. 1992; Kempe and Kazmierczak 1994; Martin-Jézéquel et al. 2000; Marron et al. 2016), it is proposed that what began as a waste-removal system may have initiated the process that resulted in biomineralization (Brennan et al. 2004). An ancestral mechanism for metal detoxification is sequestration within lysosomes (Berry et al. 1997; More et al. 2024). Indeed, one of the major proteins discovered to be important for sponge silicification is silicatein, a protein that is highly similar to members of the cathepsin L and papain family of lysosomal proteases (Shimizu et al. 1998). Similarly, cathepsins have been shown to play a role in calcification in mussels (Yarra, Ramesh, et al. 2021), sea urchins (Sciani et al. 2013), and corals (Levy et al. 2021). The V-H^+^-ATPase (VHA), a proton pump responsible for lysosomal acidification, is critical for biosilicification in diatoms, allowing for silicic acid accumulation in the SDV (Yee et al. 2020). VHA has also been linked to biomineralization in various calcifying organisms (Ziegler et al. 2004; Mackinder et al. 2011; Toyofuku et al. 2017; Iwayama et al. 2019; Hu et al. 2020; Yarra, Ramesh, et al. 2021; Achilleos et al. 2024). Interestingly, lysozymes, enzymes that can hydrolyze bacterial cell wall peptidoglycans, which have been shown to localize to lysosomes along with cathepsin D (Radons et al. 1994) play a role in avian eggshell formation (Hincke et al. 2000) and have been used for biomimetic mineralization (Luckarift et al. 2006; Wang et al. 2025). There is also transcriptional evidence that Lsi2, a silicon transporter essential for plant silicification which belongs to the same gene family as lysosome-related-organelle protein OCA2, may be involved in diatom (Shrestha et al. 2012) and sponge silicification (Maldonado et al. 2020; Francis et al. 2023; Leria and Maldonado 2026). Lsi2 genes have strong sequence orthology and functional similarity to arsenic efflux pump ArsB and are thus used interchangeably in sponge silicification studies (Marron et al. 2016; Maldonado et al. 2020; Leria and Maldonado 2026). Here, we took advantage of single-cell gene expression data, gene and protein *in situ* localization, and functional assays to visualize spicule growth in live sponges and provide novel dynamic insights about the cellular and molecular mechanisms that underlie sponge silicifcation. We show that multiple lysosome-associated genes are highly expressed in spicule forming cells called sclerocytes. Further, we demonstrate that VHA protein localizes to the sclerocyte SDV and pharmacological inhibition of VHA abrogates silica spicule formation. Using Hybridization Chain Reaction Fluorescence In Situ Hybridization (HCR-FISH), we confirmed that sponge Lsi2 genes are highly expressed in sclerocytes.

Biomineralization has evolved independently multiple times, and it was proposed to be driven by the co-option of genes within a conserved ancestral genetic toolkit (Murdock 2020): however, a core set of broadly-shared genes within the biomineralization toolkit has not been well described, and functional evidence is even more rare. Our findings indicate that lysosomal-derived machinery plays an important role in sponge biosilicification and more broadly, may have been co-opted independently multiple times for biomineralization.

## Results and Discussion

### Upregulation of lysosomal genes in sclerocytes

Analysis of single-cell RNA sequencing (scRNASeq) data of demosponge *Spongilla lacustris* (Musser et al. 2021) revealed that mRNA transcripts of genes involved in lysosomal machinery were upregulated (log2fold change > 1) in sclerocytes (Figure 1); including most subunits of VHA, an ATP-driven proton pump essential for lysosomal acidification (Mindell 2012), OCA2, also known as the pink-eyed dilution protein or P protein (H. Brilliant 2001), which localizes to lysosome related organelles called melanosomes in vertebrate pigment-producing cells (Sauka-Spengler and Bronner-Fraser 2008; Sitaram et al. 2009), lysosomal cysteine protease cathepsin-L proteins, silicateins (Shimizu et al. 1998), TMEM55A/B and TMEM192, proteins involved in lysosomal autophagy, phagocytosis, and lysosome repair (Nguyen et al. 2017; Morioka et al. 2018; Jeong et al. 2024), TMEM199, a VHA assembly factor (Miles et al. 2017), SNX8, a protein that enables lysosome reformation (Li et al. 2024), and CLIP4 and RNF11, proteins that mediate processes leading to lysosomal degradation (Mattioni et al. 2020; Tang et al. 2020).

**Figure 1.**
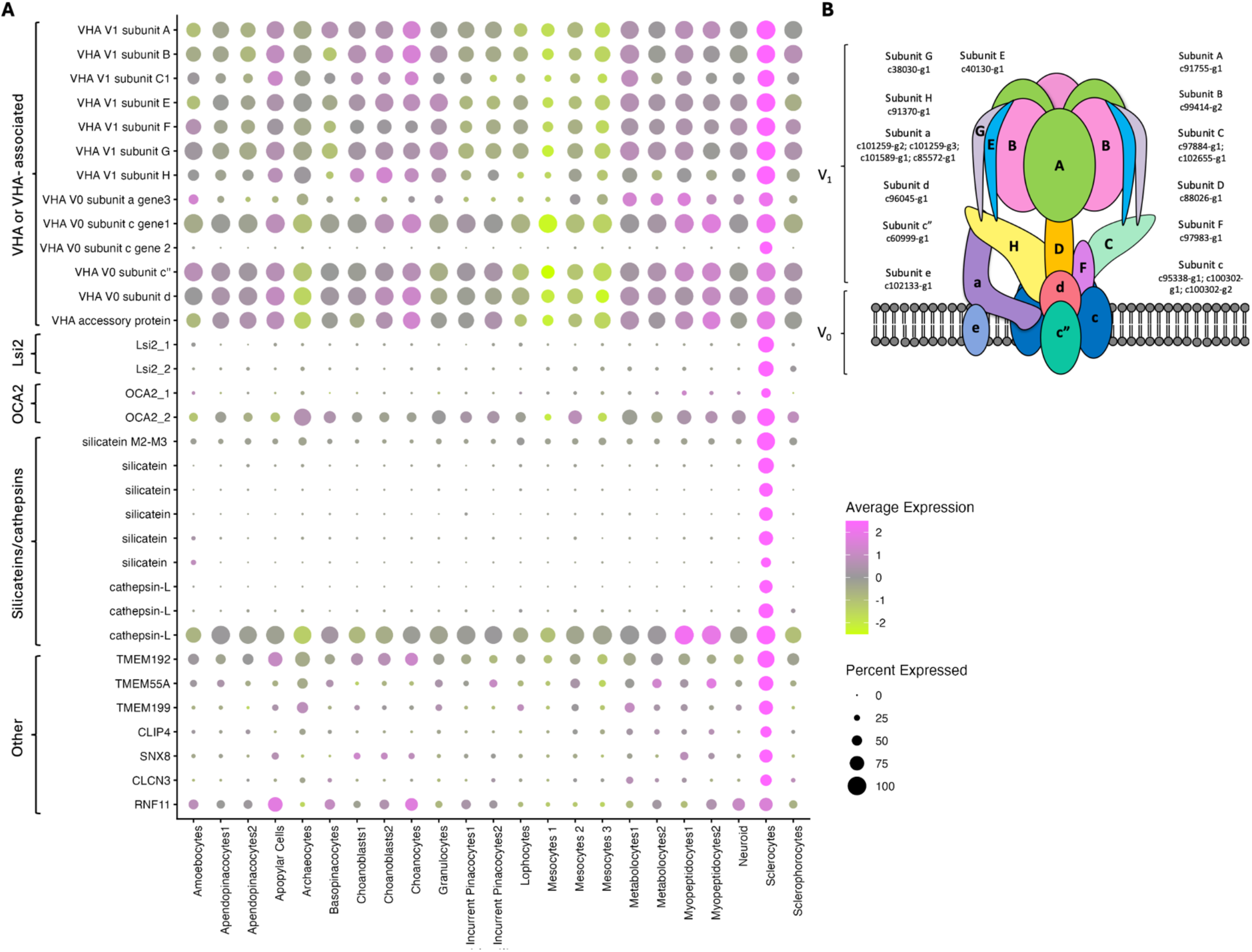
Single-cell RNA-Seq expression of lysosome-related sclerocyte markers. (A) Heatmap showing expression of genes upregulated (log2fold>1) in sclerocytes that are linked with lysosomal function across *S. lacustris* cell types (scRNASeq data from Musser et al., 2021). Note: not all VHA subunits are present here as they did not all make the cutoff log2fold cutoff. (B) Cartoon of VHA showing *S. lacustris* genes encoding each subunit.

VHA is highly conserved across all eukaryotes (Kibak et al. 1992), plays a critical role in biosilicification in diatoms (Yee et al. 2020), and has been associated with calcification in various organisms (Ziegler et al. 2004; Mackinder et al. 2011; Toyofuku et al. 2017; Iwayama et al. 2019; Hu et al. 2020; Yarra, Ramesh, et al. 2021; Achilleos et al. 2024). Further, we found that two of the upregulated sponge genes annotated as OCA2 bore greater structural resemblance to plant silicon efflux transporter Lsi2 (Figure 2) and are thus referred to as sponge Lsi2 genes in this study. However, it is worth noting that Lsi2 sponge genes are often called ArsB in the literature (Maldonado et al. 2020; Leria and Maldonado 2026). Due to the roles of VHA and Lsi2 in biomineralization in other organisms, we specifically explored their roles in sponge biosilicification.

**Figure 2.**
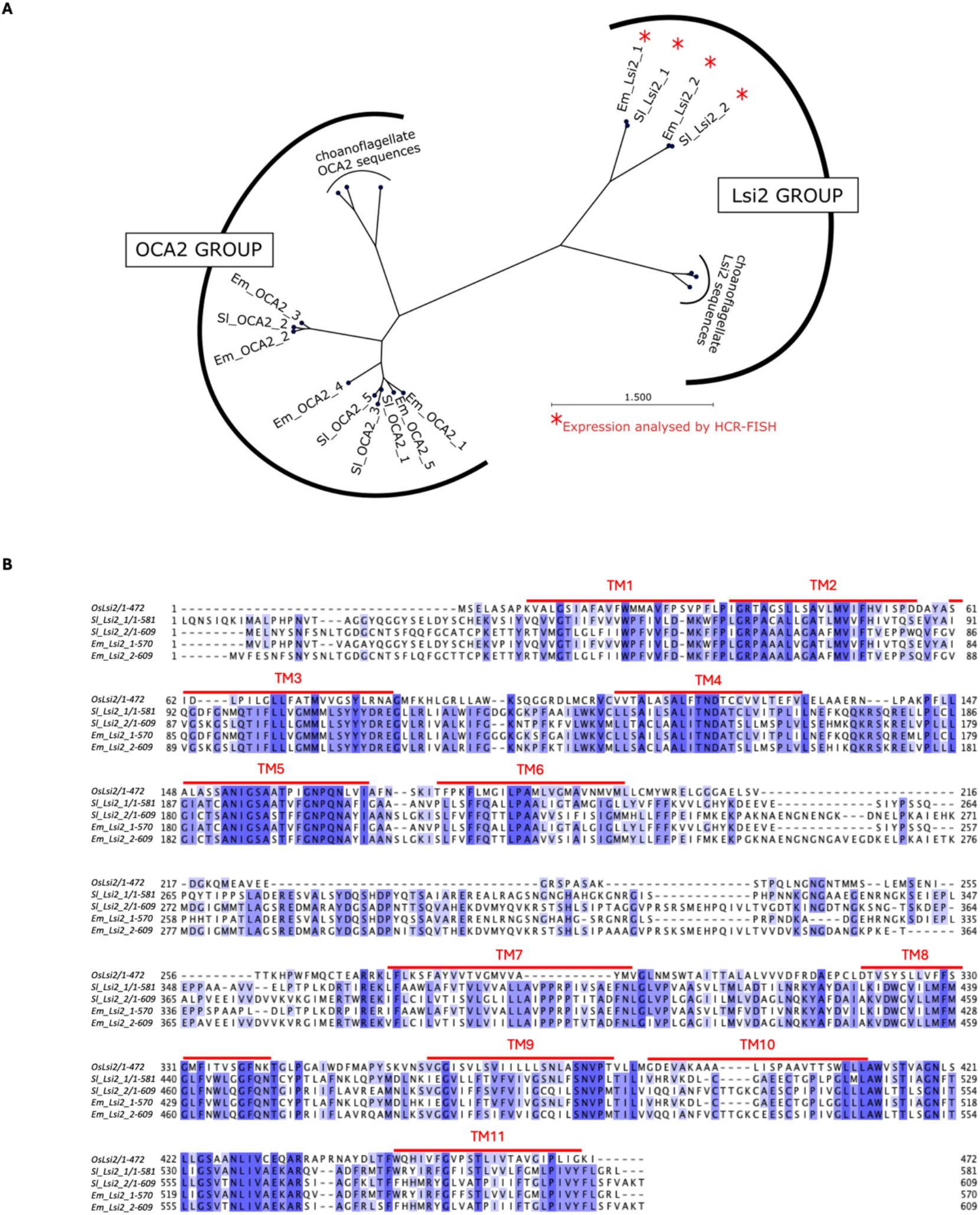
Phylogenetic and structural analysis of Lsi2/OCA2 homologs. (A) Maximum-likelihood phylogeny of Lsi2 and OCA2 homologs in *Ephydatia muelleri* (abbreviated Em) and *Spongilla lacustris* (abbreviated Sl). (B) Sequence alignment of *OsLsi2* gene from rice (*Oryza sativa)*, Lsi2_1 and 2 from *E muelleri*, and Lsi2_1 and 2 from *S. lacustris*. Alignment shaded from light blue to dark blue based on percentage identity similarity. 11 predicted transmembrane (TM) domains in red (based on Ma et al., 2007).

### VHA is abundant in sclerocytes and is essential for skeleton formation

VHA plays a wide range of physiological roles in marine organisms (Tresguerres et al. 2007; Tresguerres et al. 2013; Barott et al. 2015; Tresguerres 2016; Yee et al. 2020; Barott et al. 2022; Hargadon et al. 2024; Allard et al. 2025; Thies et al. 2025). Notably, VHA is essential for biosilicification in diatoms, allowing for silicic acid accumulation in the SDV (Yee et al. 2020). VHA has been linked to biomineralization in calcifying organisms as well. In foraminifera, VHA maintains the proton flux responsible for calcification (Toyofuku et al. 2017). The c/c′ subunit of the V0 sector of VHA has been shown to be upregulated in calcifying strains of coccolithophores (Mackinder et al. 2011). A transcriptomic study on bryozoans identified VHA as a candidate biomineralization protein as it was upregulated in younger zooids, which are at the distal growth margin of the colony, compared to older zooids (Achilleos et al. 2024). Similarly, VHA was transcriptionally identified to be involved in bivalve shell remodeling (Yarra, Blaxter, et al. 2021). In sea urchin larvae, VHA has been shown to play an essential role in skeleton re-calcification (Hu et al. 2020). VHA also contributes to the mineralization-demineralization cycle of terrestrial isopods (Ziegler et al. 2004).

Our analysis of published scRNASeq data from *S. lacustris* (Musser et al. 2021) found that all subunits of VHA were upregulated in sclerocytes compared to other cell types, with the exception of integral domain V0 subunit e which was more highly expressed in myopeptidocytes (Supplementary Figure 1). However, this subunit does not seem to be essential for proton pump activity and may instead play a role in endomembrane specific VHA-assembly or targeting (Seidel et al. 2008). Some subunit isoforms such as V0 subunit a gene 3 and V0 subunit c gene 2 were specifically upregulated in sclerocytes compared to other cell types and to other isoforms suggesting that these isoforms may only localize to sclerocyte-specific organelles, presumably the SDV. Indeed, immunostaining with specific antibodies against VHA subunit B (VHA_B_) revealed a striking VHA localization within sclerocytes and around spicules in *S. lacustris* as well as in *Ephydatia muelleri* (Figure 3), another freshwater sponge. In both cases, the VHA protein appeared to be surrounding the SDV within sclerocytes, and was distributed throughout the cytoplasm, potentially on vesicles (Supplementary Figure 2). Airyscan ‘super-resolution’ imaging showed VHA surrounding spicules fluorescently labelled with PDMPO [2-(4-pyridyl)-5-((4-(2-dimethylaminoethylaminocarbamoyl)methoxy)-phenyl)oxazole], an acidotropic dye that is selectively incorporated and co-deposited with Si into the newly polymerized silica (27) (Figure 4A). To test the functional role of VHA in spiculogenesis, sponges were incubated with 100 nM of the specific pharmacological inhibitor of VHA, concanamycin (concA) (Dröse and Altendorf 1997; Huss et al. 2002), for 24 hours along with PDMPO or with vehicle DMSO as a control (Figure 4B). VHA inhibition significantly decreased the number of spicules per mm^2^ of actively growing sponge tissue by 41.6% ± 10.6% in *E. muelleri* and 77.5% ± 7.1% in *S. lacustris*. The proportion of spicules labeled with PDMPO decreased by 85.8% ± 4.7% in *E. muelleri* and 98.3%± 1.7% in *S. lacustris* indicating that spiculogenesis is stunted by VHA inhibition (Figure 4C-D). Given that *S. lacustris* and *E. muelleri* are phylogenetically distinct sponges with divergent life history, these observations suggest that VHA-dependent silicification is a conserved mechanism in freshwater demosponges.

**Figure 3.**
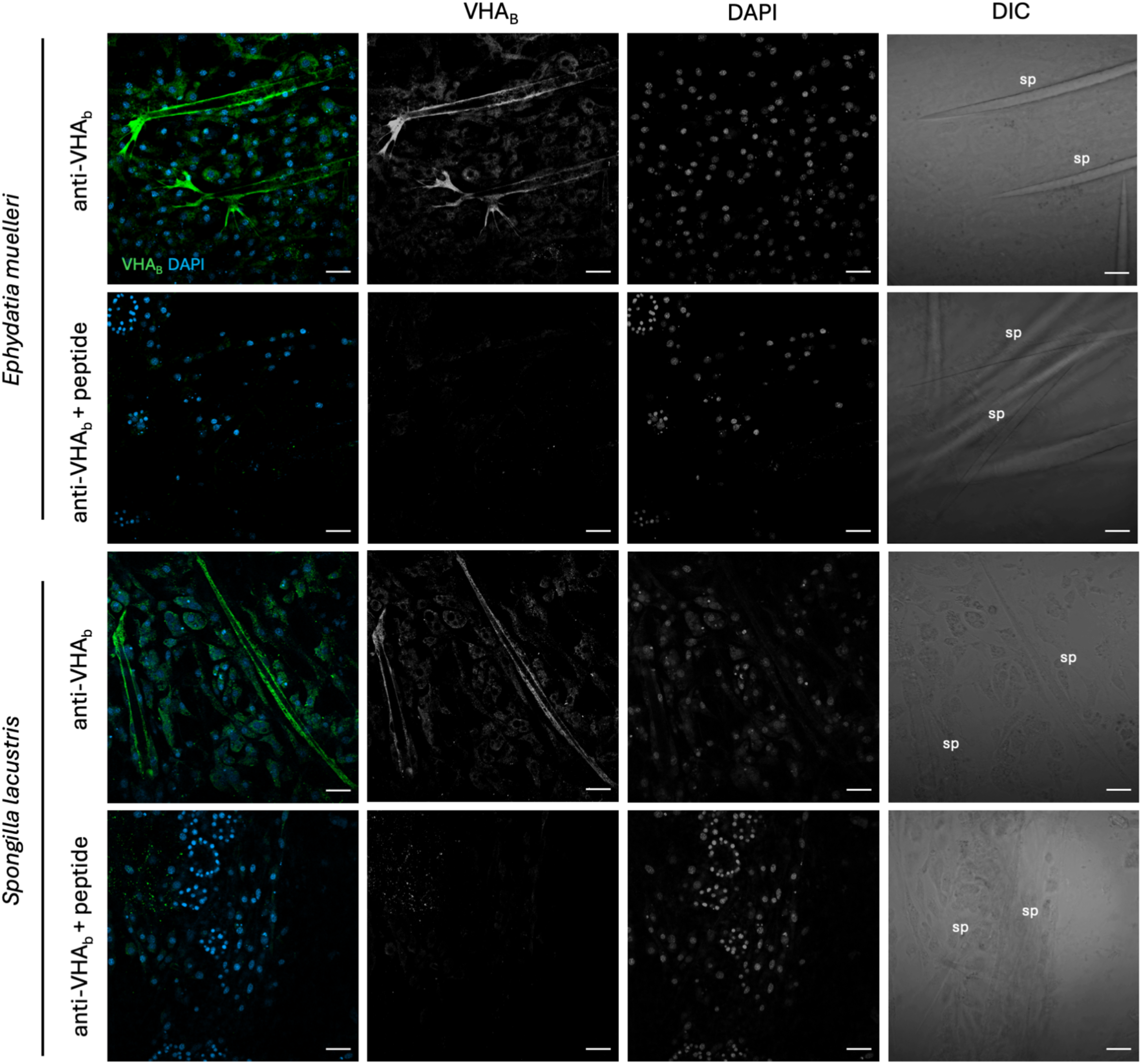
VHA localization in *Ephydatia muelleri* and *Spongilla lacustris*. Immunohistochemical localization of VHA_B_ in *E. muelleri* and *S. lacustris* sponges incubated with anti-VHA_B_ antibody or anti-VHA_B_ antibody with excess peptide (peptide pre-absorption control). VHA_B_ in green, DAPI in blue. VHA_B,_ DAPI, and DIC channels are also shown separately. All images except DIC are orthogonal projections. Scale bars: 20 μm. sp: spicules.

**Figure 4.**
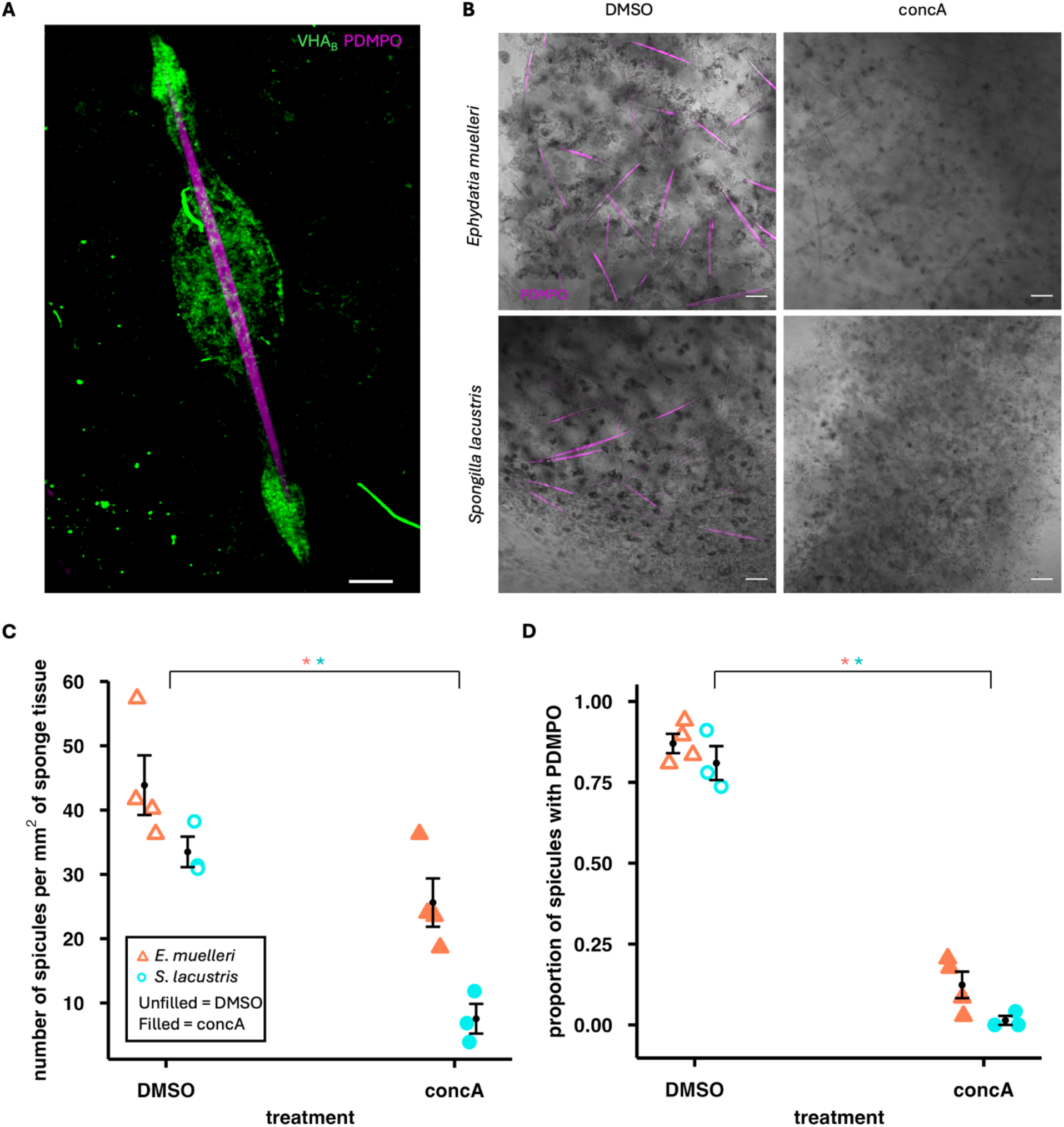
Essential role of VHA in spiculogenesis in *Ephydatia muelleri* and *Spongilla lacustris*. (A) Airyscan confocal image showing VHA_B_ localization around *E. muelleri* spicule labelled with PDMPO (B) Representative images of PDMPO fluorescence of spicules in sponges incubated with DMSO or 100 nM concA for 24 hours. (C) Dot plot depicting the number of spicules per mm^2^ of sponge tissue incubated with DMSO or 100 nM concA for 24 hours with error bars. (D) Dot plot depicting average proportion of spicules that showed PDMPO fluorescence in sponges incubated with DMSO or 100 nM concA for 24 hours with error bars. *E. muelleri* in red (n=4) and *S. lacustris* in cyan (n=3). Asterisks denote statistical significance (t test, p-value < 0.05). VHA_B_ in green, PDMPO in magenta. Scale bars in (A): 5 μm and (B): 50 μm.

These results show that, as reported in diatoms (Yee et al. 2020), VHA-dependent SDV acidification is critical for sponge biosilicification. By acidifying the SDV in both diatoms and sclerocytes, VHA drives silicic acid accumulation and prevents its rapid polymerization into silica (Vrieling et al. 1999; Mejía et al. 2013). Given prior evidence that VHA plays a role in calcification across a wide range of phyla, VHA appears to have been co-opted independently by multiple lineages for biomineralization. Interestingly, plants also use a proton pump to acidify the xylem sap where silicic acid is stored prior to hardening. The xylem is acidified by the plant and fungi-specific (Baxter et al. 2005) plasma membrane P-type H^+^-ATPase (Jahn et al. 1998; Sattelmacher 2001). The co-option of proton pumps for maintenance of accumulated silicic acid appears to be a shared feature for biosilicification over vastly diverged lineages.

### Lsi2 and OCA2 are highly expressed in sclerocytes

Lsi2 and OCA2 belong to a family of transmembrane transporters with similarity to SLC13 (Kondo et al. 2015; Marron et al. 2016; Maldonado et al. 2020; Coskun et al. 2021; Mesdaghi et al. 2023). Lsi2 is a silicic acid/H^+^ antiporter that mediates silicic acid efflux into the plant xylem (Ma et al. 2007). OCA2 is an anion channel that localizes to melanosomes, lysosome related organelles, in pigment-producing cells called melanocytes (Sitaram et al. 2009) which specifically occur in vertebrates (Sauka-Spengler and Bronner-Fraser 2008). OCA2mediates chloride-selective anion conductance, providing charge compensation and increasing proton-pumping efficiency of VHA (Stauber and Jentsch 2013; Bellono et al. 2014). A phylogenetic analysis of the evolutionary history of Si transport reported that Lsi2 has homologs across a broad range of eukaryotes which resolve into two main groups: one containing vertebrate OCA2-like sequences, and the other containing the land plant Lsi2s together with Lsi2-like genes from diverse prokaryotic and eukaryotic organisms (Marron et al. 2016). This Lsi2-like clade was dominated by homologs from silicifying organisms such as land plants, sponges, loricate choanoflagellates, haptophytes, chrysophytes/synurophytes, radiolarians, and diatoms indicating that Lsi2-like genes play a role in biosilicification. In alignment with this, Lsi2-like genes were upregulated in marine hexactinellid sponge *Vazella pourtalesii* upon exposure to dissolved silicon (dSi) (Maldonado et al. 2020) and a detailed phylogenetic analysis showed that these genes were present in ancestral sponges and were retained and expanded in silicifying sponge lineages (Leria and Maldonado 2026). In another marine hexactinellid sponge *Aphrocallistes vastus*, an analysis of differentially expressed genes between the osculum, the vertical growth zone where new skeletal growth is concentrated (Kahn and Leys 2017), and the main body, reported that Lsi2 was upregulated with other mineralization-related genes (Francis et al. 2023). Finally, transcriptomic evidence suggests that an Lsi2-like gene may be involved in silicic acid transport in diatoms (Shrestha et al. 2012). However, the cellular localization of the expressed genes has not been reported in the literature.

Our analysis of published scRNASeq data from *S. lacustris* (Musser et al. 2021) showed that Lsi2 genes were upregulated in sclerocytes (Figure 5A). Alignment of the protein sequences of the Lsi2 genes in *S. lacustris* and their equivalent homologs in *E. muelleri* against rice *Oryza sativa* Lsi2 (*Os*Lsi2) showed that transmembrane regions are highly conserved (Figure 2B). OCA2 genes were also upregulated in sclerocytes but OCA2_1 was expressed in a smaller percentage of sclerocyte cells than the Lsi2 genes and OCA2_2 was expressed in other cell types as well indicating it may play a more generic housekeeping role, possibly in sponge lysosomes within all cells. Sponge OCA2 genes may play a role similar to their function in vertebrate melanocytes of providing counter-ion balance and improving efficiency of the VHA acidification mechanism. Notably, we found that Lsi2 gene expression was much higher in sclerocytes than the previously described sponge silicic acid uptake protein the {Na^+^/HCO_3_^−^[Si(OH)_4_]} co-transporter or NBCSA (Schröder et al. 2004) (54A) which matches recent claims that NBCSA is not critical for silicic acid transport in all sponges (Maldonado et al. 2020). Confirming the scRNASeq data, HCR-FISH probes designed against the Lsi2 genes in *S. lacustris* and *E. muelleri* distinctly labeled sclerocytes (Figure 5B). Interestingly, the two Lsi2 homologs showed heterogeneous expression in sclerocytes: i.e. Lsi2_1 signal was stronger in some sclerocytes, while Lsi2_2 signal was stronger in others; overall, Lsi2_2 was more ubiquitously expressed around spicules than Lsi2_1 (Supplementary Figure 3). As we only visualized mRNA, we could not definitively conclude whether the Lsi2 proteins localize to the SDV membrane or the plasma membrane. Unfortunately, our attempt to design antibodies against these proteins were unsuccessful. However, we speculate that they would localize to the SDV membrane as the Lsi2 silicic acid/H^+^ antiporter activity relies on a proton gradient and VHA would only be able to pump H^+^ ions from the cytosol to an intracellular compartment (Forgac 2007). Our results provide novel transcriptomic and spatial evidence to support previous claims of the role of Lsi2 in sponge silicic acid uptake (Marron et al. 2016; Maldonado et al. 2020; Leria and Maldonado 2026).

**Figure 5.**
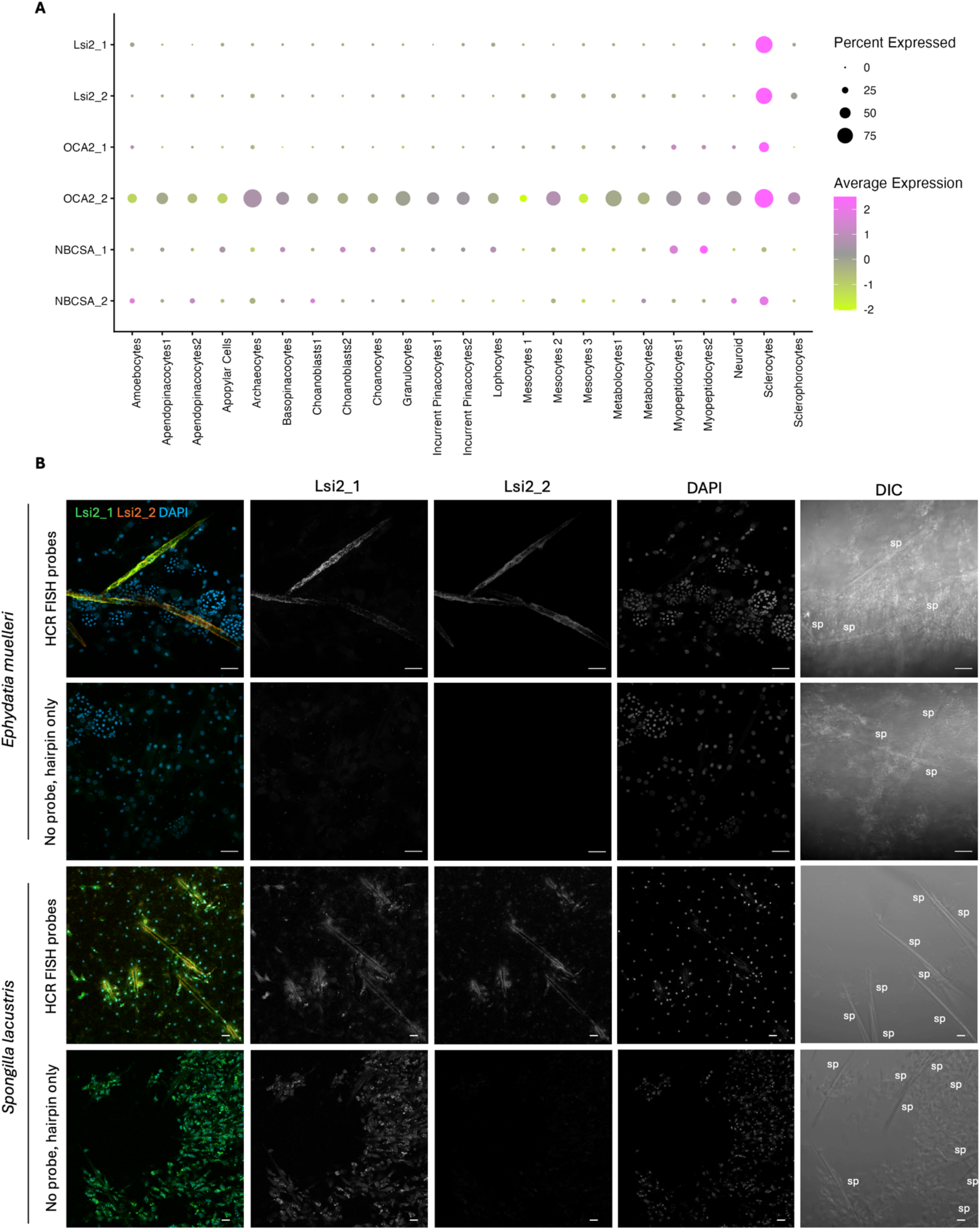
Gene expression of Lsi2 genes in *Ephydatia muelleri* and *Spongilla lacustris*. (A) Heatmap showing expression of Lsi2, OCA2, and [Na^+^/HCO_3_^-^[Si(OH)_4_]] co-transporter (NBCSA) genes in *S. lacustris* cell types. (B) HCR-FISH in *E. muelleri* and *S. lacustris* sponges incubated with probes designed against Lsi2_1 and Lsi2_2 genes. Lsi2_1 in green, Lsi2_2 in orange, DAPI in blue. Lsi2_1, Lsi2_2, DAPI, and DIC channels are also shown separately in grayscale. All images are orthogonal projections. Scale bars: 20 μm. sp: spicules

In addition to their findings on Lsi2, Maldonado identified a homolog to passive plant silicic acid transporter Lsi1 called the aquaglyceroporin channel (gAQP3/9) which was upregulated in *V. pourtalesii* upon exposure to dSi (Maldonado et al. 2020) and appears to have coevolved with Lsi2 in multiple lineages (Leria and Maldonado 2026). We did not find significant upregulation of homologs of Lsi1/gAQP3/9 in sclerocytes compared to other cell types in our scRNASeq analysis (Supplementary Figure 4), but this does not rule out their involvement in silicification as the genes are positively expressed in sclerocytes and it is very likely that they are involved in multiple other cellular functions. Leria and Maldonado 2026 hypothesize that, in addition to sclerocytes, Lsi1/gAQP and Lsi2 transporters are localized to pinacocytes to facilitate transport of dSi into the sponge from the external environment. While we see evidence of gAQP homologs in apendopinacocytes and incurrent pinacocytes, they are once again not significantly upregulated compared to other cell types (Supplementary Figure 4). Meanwhile, we see very minimal expression of Lsi2 homologs in pinacocytes in the scRNASeq data and in our HCR FISH localization studies (Figure 5). These findings indicate the Lsi2 and Lsi1/gAQP3/9 may not be used in pinacocytes for silicic acid uptake. Instead, an alternative uptake mechanism may be at play, or sclerocytes might directly take up silicic acid from the environment. These results highlight the importance of cell type specific transcriptomic and spatial resolution for deciphering complex mechanisms.

### Links between lysosomal detoxification and biomineralization

In their phylogenetic analysis of silicon transport genes, Marron *et al*. propose that the Lsi2 protein evolved in the original eukaryote as a detoxification mechanism to export silicic acid out of cells in the high Si oceans of the Precambrian to prevent uncontrolled silica deposition and were later co-opted for biosilicification by a specific set of lineages at the Precambrian/Cambrian boundary (Marron et al. 2016). This hypothesis implies a switch of Lsi2 orientation within the membrane from exporting silicic acid out of cells to transporting it in. Alternatively, it is possible that Lsi2 detoxified silicic acid by sequestering it within lysosomes rather than by exporting it out of cells. Such a mechanism would seamlessly lend itself to the accumulation of silicic acid into the SDV of diatoms and sponges, while also inserting at the plasma membrane of plant cells in an orientation that mediates silicic acid extrusion into the xylem.

Lysosomes and detoxification are intrinsically linked. For example, calcium and heavy metal ions including zinc, copper, and iron are sequestered in lysosomes and vacuoles in many invertebrate phyla into structures termed concretions or granules, rendering them insoluble and detoxified (Howard et al. 1981; Engel 1983; Ballan-Dufrançais 2002; Wallace et al. 2003; Desouky 2006; Pigino et al. 2006; Ahearn et al. 2010; Wang et al. 2011; Marin 2012). These organelles are often acidified by VHA (Sterling et al. 2007; Blaby-Haas and Merchant 2014). The sequestered concretions found in oysters (Wang et al. 2011), bivalves (Wallace et al. 2003), snails (Howard et al. 1981; Desouky 2006) insects (Ballan-Dufrançais 2002), and crustaceans (Sterling et al. 2007; Ahearn et al. 2010) could be considered precursors to skeletons. In fact, detoxification was likely a selective pressure that shaped the evolution of biomineralization (Simkiss 1977; Carney et al. 2007; Li et al. 2022). Calcium ions are disruptive to cellular functioning as their precipitation intracellularly would be deleterious; therefore, sequestration within a pre-existing acidified organelle may have provided an energetically effective solution (Simkiss 1977), while also offering a parsimonious mechanistic explanation at the evolutionary level. Recently, it was shown that mouse osteoblasts, cells responsible for secreting calcium phosphate mineral precursors for bone formation, transport amorphous calcium phosphate via lysosomes (Iwayama et al. 2019). Additionally, pharmacological inhibition of VHA prevented the formation of electron-dense intracellular granules and extracellular calcified nodules, (Iwayama et al. 2019) matching the importance of VHA for silicification in sponges (current study) and in diatoms. (Yee et al. 2020) Furthermore, other studies have shown that increased expression of the master regulator of lysosomal biogenesis, *Tfeb*, enhances osteoblastic differentiation (Tsukuba et al. 2017) and increases bone mass and strength (James et al. 2025). Altogether, the evidence suggests that ancient links between detoxification and lysosomal machinery set the stage for the repeated independent evolution of biomineralization. However, this does not rule out other possibilities for the co-option of lysosomal genes for skeleton formation. For example, lysosomes are also essential for microbial defense (Levine and Deretic 2007). The role of anti-bacterial protein lysozyme in avian eggshell formation (Hincke et al. 2000) might indicate that interactions with microbes also played a role in the emergence of biomineralization in some species.

### Lysosomal machinery as part of ancestral genetic toolkit for biomineralization

It has been proposed that genes involved in biomineralization were independently co-opted from an ancestral genetic toolkit (Murdock 2020), however, the specific genes within this toolkit have not been clearly identified. Here, we found high expression of lysosomal-associated genes in sclerocytes including cathepsins, silicateins, TMEM55A/B, TMEM192, and TMEM199. Cathepsins are lysosomal proteases speculated to play a role in making precise cuts in matrix and mineralization proteins for the process of biosilicification (Francis et al. 2023). They are upregulated during shell repair in mussels (Yarra, Ramesh, et al. 2021), cathepsin B/X has been identified in sea urchin spine extracts (Sciani et al. 2013), and our analysis of scRNASeq dataset of a stony coral revealed that cathepsin L and cathepsin V are upregulated in skeleton-forming cells, calicoblasts (Levy et al. 2021). Silicatein was discovered in the protein filaments of spicules from demosponge *Tethya aurantia* (Shimizu et al. 1998). Silicateins are highly similar to members of the cathepsin L and papain family of proteases, but the cysteine at the active site in the proteases is replaced by serine (Shimizu et al. 1998). They have been proposed to retain protease activity similar to other cathepsins (Müller et al. 2008), catalyze the polymerization of silicic acid (Povarova et al. 2018), or serve as scaffolding for spicule formation (Görlich et al. 2020). TMEM55A/B and TMEM192, involved in lysosomal autophagy, phagocytosis, and lysosome repair (Nguyen et al. 2017; Morioka et al. 2018; Jeong et al. 2024), and TMEM199 (Miles et al. 2017), a VHA assembly factor, have not been previously described within the context of biosilicification to our knowledge. We found upregulation of VHA mRNA in sclerocytes, VHA protein localization around spicules, and showed that inhibition of VHA disrupts skeleton formation. This aligns with similar findings on biosilicification in diatoms (Yee et al. 2020), and calcification in coccolitophores (Mackinder et al. 2011), foraminifera (Toyofuku et al. 2017), sea urchins (Hu et al. 2020), bryozoans (Achilleos et al. 2024), and crustaceans (Ziegler et al. 2004). We report that Lsi2 genes were upregulated in sponge sclerocytes, likely playing a role in silicic acid uptake similar to plants (Ma et al. 2007). In the marine hexactinellid sponge *Aphrocallistes vastus*, VHA, Lsi2, and various cathepsins were upregulated in the osculum region, indicating a potential link to sponge skeleton formation (Francis et al. 2023). Exposure of hexactinellid sponge *Vazella pourtalesii* to dSi, led to the upregulation of cathepsins and Lsi2 (Maldonado et al. 2020). Further, a proteomic analysis found that spicule contents of marine demosponge *Petrosia ficiformis* were dominated by proteins involved in lysosomal function including cathepsins and silicatein (Pozzolini et al. 2022). A recent phylogenetic study found that biomineralization evolved independently in sponges multiple times (Rossi et al. 2026). Notably siliceous spicules seem to have evolved once in Hexactinellida, twice in Demospongiae, and once in Homoscleromorpha (Rossi et al. 2026). These different lineages evolved differing strategies for silica polymerization; hexactinellids use hexaxilin, glassin, and perisilin protein to polycondense silicic acid into spicules (Shimizu et al. 2015; Shimizu et al. 2024) demosponges use silicatein (Shimizu et al. 1998; Müller et al. 2007; Mohri et al. 2008), and the silicifying proteins of sponges in the class Homoscleromorpha remain unidentified (Leria and Maldonado 2026). However, past transcriptomic data corroborated by our spatial and functional work indicate that VHA and Lsi2, the mechanisms for SDV acidification and silicic acid uptake, were ancestral to sponges and thus must have been independently co-opted multiple times as biosilicification emerged.

While the many studies linking biomineralization and lysosomal genes largely relied on methods like bulk RNASeq, phylogenetics, and proteomics of isolated spicules, a major advance of our study is the usage of scRNASeq data, *in situ* protein and mRNA localization, and functional assays to conclusively show the importance of lysosomal genes in sponge skeleton formation. The machinery responsible for the biogenesis of lysosome-related organelles is present across the eukaryotic tree of life and is likely to have been in place prior to the last eukaryotic common ancestor (More et al. 2024). Given that lysosomal genes have been shown to be involved in biomineralization across a broad range of taxa, the combined evidence suggest that lysosomal machinery may have been co-opted independently multiple times for skeleton formation, forming a key set of genes within the ancestral toolkit for biomineralization.

## Conclusion

Skeleton formation was a major innovation that catapulted the trajectory of body plan evolution during the Cambrian. It provided organisms with new possibilities for protection, support, locomotion, and defense (Gilbert et al. 2022). Biomineralization is intrinsically linked with global calcium and silica cycling. Importantly, many marine organisms are ballasted by their silica and calcium skeletons for burial in the seafloor, allowing for carbon sequestration (Armstrong et al. 2001; Aldridge et al. 2012). Here, we describe the co-option of lysosomal genes for biosilicification in sponges, the earliest diverged branch of metazoans (Steenwyk and King 2025). We discuss parallels to biomineralization pathways across the tree of life, from protists to mammals. Although biomineralization evolved independently multiple times, the co-option of genes from a conserved toolkit is remarkable (Murdock 2020). Our findings, combined with data and ideas from previous studies, provide compelling evidence for the co-option of ancestral lysosomal machinery for biosilicification, and potentially for biomineralization in general.

## Materials and Methods

### Organisms

*Ephydatia muelleri* and *Spongilla lacustris* gemmules were obtained from the Nichols lab. They were collected from the upper peninsula of Michigan in 2023. Gemmules were stored at 4°C and hatched in spring water (Crystal Geyser) at room temperature in cover-slip bottom dishes as previously described (Nichols 2023a).

### scRNASeq analysis

Datasets acquired from Musser et al. (Musser et al. 2021) and used Seurat in R to plot the expression of sclerocytes markers. We referenced the *S. lacustris* transcriptome and proteome provided in Musser et al. (Musser et al. 2021) to identify homologs and design HCR-FISH probes. R code and lists of *S. lacustris* genes analyzed can be found in our [github repository](https://github.com/maitripaul/Co-option-of-Lysosomal-Machinery-for-Biomineralization/tree/main).

### Phylogenetic and structural analysis of Lsi2/OCA2 homologs

Lsi2 and OCA2 homologs were identified in *Ephydatia muelleri* (abbreviated Em) and *Spongilla lacustris* (abbreviated Sl) genomic and transcriptomic datasets by BLAST searches using choanoflagellate sequences from Marron *et al*.(Marron et al. 2016) and by extraction of candidate orthogroups from OrthoFinder. Protein sequences were aligned with MAFFT using the DASH structure-guided alignment mode, after which DASH profile sequences were removed. The alignment was trimmed with ClipKIT to retain phylogenetically informative sites, and a maximum-likelihood phylogeny was inferred using IQ-TREE with automated model selection and branch support assessed by 1,000 ultrafast bootstraps and 1,000 SH-aLRT replicates.

Sequence alignments of *OsLsi2* gene from rice (*Oryza sativa)*, Lsi2_1 and 2 from *E muelleri*, and Lsi2_1 and 2 from *S. lacustris* were done on Clustal Omega and visualized using Jalview.

### Antibodies

Sponges were probed for VHA using a custom rabbit polyclonal antibody developed using a peptide antigen matching a conserved region of VHA’s B subunit (VHA_B_; AREEVPGRRGFPGY; GenScript Biotech Corporation, Piscataway, NJ, USA) which is conserved in species ranging from sponges to humans (Tresguerres et al. 2013; Barott et al. 2015; Roa and Tresguerres 2016; Armstrong et al. 2018; Damsgaard et al. 2020). We confirmed the epitope is 100% conserved in *E. muelleri* and *S. lacustris* (Supplementary Figure 5A).

### Immunohistochemistry

Immunohistochemical localization of anti-VHA_B_ antibody in sponges was done following previously described protocol (Nichols 2023b). Briefly, 4-7 day old gemmule-hatched *E. muelleri* and *S. Lacustris* sponges were fixed with 4% paraformaldehyde in 95% EtOH for 45 minutes at room temperature. Sponges were rinsed with 0.1% Tween 20 (v/v) in phosphate buffered saline (PTw) three times before incubation with blocking buffer (3% Bovine Albumin Serum in PTw) for 45 minutes at room temperature. Samples were incubated overnight (4°C) with either 0.962 μg ml^−1^ polyclonal anti-VHA_B_ primary antibody in blocking buffer or 0.962 μg ml^−1^ polyclonal anti-VHA_B_ primary antibody with 60x peptide as a “peptide preabsorption control”. Sponges were rinsed with PTw three times and subsequently incubated with 4 μg ml^−1^ secondary antibody (goat anti–rabbit–Alexa Fluor 488; A-11008; Invitrogen, Carlsbad, CA, USA) and 4 μg ml^−1^ 4′,6-diamidino-2-phenylindole (DAPI) DNA stain (Invitrogen) in blocking buffer for 45 minutes in the dark at room temperature. Excess secondary antibody and DAPI were washed from samples by rinsing three times with PTw and they were mounted for imaging using ProLong Glass Antifade Mountant (P36982; Invitrogen). Confocal microscopy was performed on a Zeiss AxioObserver Z1 connected to a laser scanner equipped with 405, 488, 561, and 640 nm laser lines (Zeiss LSM 800 with Airyscan, Carl Zeiss AG, Oberkochen, Baden-Württemberg, Germany). Samples were imaged with the following excitation and emission wavelengths: Alexa Fluor 488 — ex: 488 nm, em: 488-561 nm; DAPI—ex: 405 nm, em: 400-488 nm.

### Western blots

Approximately 50-80 *E. muelleri* and *S. lacustris* gemmules were hatched in separate petri dishes. At 9-11 days old, sponges were scraped off the dishes with a razor blade and dounce homogenized in 100 μL of 1x protease inhibitor cocktail (P2714; Sigma-Aldrich) in spring water. Samples were sonicated for 3x10 second bursts with 1-minute rests on ice. Homogenates were spun down for two minutes and the supernatant was retained. The protein concentrations of samples were obtained using Bradford assay performed with a bovine serum albumin standard curve (BioRad, Hercules, CA, USA). Samples were incubated in Laemmli Sample buffer with 5% (v/v) β-mercaptoethanol for 5 min at 100 °C and increasing amounts of protein were loaded into a sodium dodecyl sulphate–polyacrylamide gel electrophoresis gel in duplicate (*E. muelleri*: 0.7 μg, 3.5 μg, 7 μg, 13.9 μg; *S. lacustris*: 0.7 μg, 3.5 μg, 7 μg, 13.9 μg). The gel was run in in Tris-glycine running buffer (25 mM Tris-base, 192 mM glycine, 0.1% (w/v) sodium dodecyl sulphate, pH 8.3; Sigma-Aldrich) at 47V for 35 minutes followed by 150V for 35-40 minutes until the proteins had reached the bottom of the gel. Proteins were transferred from the gel onto a polyvinylidene difluoride (PVDF) membrane using a Mini Tans-Blot Cell (Bio-Rad) filled with Towbin transfer buffer (25 mM Tris, 192 mM glycine, 20% (v/v) methanol, pH 8.3; Sigma-Aldrich) at 90 mAmps overnight. PVDF membranes were incubated with blocking buffer (1g non-fat dry milk powder in 10 mls Tris-buffered saline with 0.1% Tween (TBS-T)) for 1 hour at room temperature on an orbital shaker and subsequently incubated overnight at 4 °C with either 0.2405 μg/mL anti-VHA_B_ primary antibody or anti-VHA_B_ primary antibody with 60x excess peptide (“peptide preabsorption control”) in blocking buffer. Membranes were washed with 3 x 10 min TBS-T and then incubated with secondary antibody (goat anti–rabbit–horseradish peroxidase diluted 1:5000; Bio-Rad) for 1 hour on an orbital shaker at room temperature. Membranes were washed again with 3 x 10 min TBS-T and imaged with Clarity Max Western ECL Substrate (Bio-Rad) using a Chemidoc Imaging system (Bio-Rad). Western blotting specifically detected the expected ∼55 kDa band (Supplementary Figure 5B).

### HCR-FISH protocol

Probes were designed using publicly available HCR-FISH probe generator (Choi et al. 2018; Tsuneoka and Funato 2020; Kuehn et al. 2022; Wang et al. 2024). Sequences against which probes were designed are provided in Supplementary Table 1 in our github repository. *E. muelleri* Lsi2 homologs were identified by blasting Lsi2 sequences against an unpublished transcriptome. 6-10 day old gemmule-hatched *E. muelleri* and *S. Lacustris* sponges were stained using HCR FISH probes following an adapted previously described protocol (Nichols 2023c). Briefly, sponges were fixed in cold 4% PFA in 1/4x HF buffer (147.5mM NaCl, 0.1675 mM KCl, 0.19 mM CaCl_2_, 0.06mM NaHCO in RNAse-free water) on a gently rocking platform at 4C overnight. Samples were rinsed with ¼ x HF buffer and then incubated with ¼ x HF buffer containing an RNAse inhibitor (1 U/μl final concentration) at room temperature for 10 minutes. Sponges were rinsed again with ¼ x HF buffer and progressively dehydrated into 100% methanol. Subsequently, 100% methanol was replaced with 100% ethanol, sponges were progressively rehydrated into 0.1% Tween 20 (v/v) in phosphate buffered saline (PTw) and incubated with of 5 μg/mL Proteinase K in PTw for 90 seconds at room temperature. The proteinase K solution was removed and sponges were incubated with 2mg/ml glycine in PTw for 5 minutes at room temperature twice. Samples were rinsed with PTw and re-fixed with 4% PFA in PTw for one hour at room temperature. Following two rinses with PTw, samples were incubated with 2x Saline Sodium Citrate (SSC) for 10 minutes at room temperature twice. Sponges were then pre-hybridized by incubation with Hybridization buffer (Molecular Instruments), first at room temperature for 10 minutes and then at 37°C for 30 minutes. Subsequently, samples were incubated with 0.04 μM probe in hybridization buffer or just hybridization buffer for the “no probe, hairpin only control” (Supplementary figure 6) at 37°C overnight. Sponges were rinsed with probe wash buffer (Molecular Instruments) for 5 mins at 37°C three times, then incubated with 2x SSC for 5 minutes at room temperature twice. Samples were pre-amplified with amplification buffer (Molecular Instruments) for 5 minutes at room temperature twice, then incubated with 60 nM of snapcooled (incubated at 95°C for 90 seconds, then cooled in the dark for at least 30 minutes) hairpins (Molecular Instruments) in amplification buffer at room temperature in the dark overnight. Specimens were rinsed with 2x SSC for 5 minutes at room temperature twice. 2x SSC was replaced with 10 μg ml^−1^ 4′,6-diamidino-2-phenylindole (DAPI) in PBS for 20 minutes at room temperature. Sponges were rinsed with PBS and then mounted using ProLong Glass Antifade Mountant (P36982; Invitrogen). Confocal microscopy was performed on a Zeiss AxioObserver Z1 connected to a laser scanner equipped with 405, 488, 561, and 640 nm laser lines (Zeiss LSM 800 with Airyscan, Carl Zeiss AG, Oberkochen, Baden-Württemberg, Germany). Samples were imaged with the following excitation and emission wavelengths: Alexa Fluor 555 — ex: 561 nm, em: 561-585 nm; Alexa Fluor 488 — ex: 488 nm, em: 488-561 nm; DAPI—ex: 405 nm, em: 400-488 nm.

### PDMPO experiments

4-11 day old gemmule-hatched *E. muelleri* and *S. Lacustris* sponges were incubated with 1 uM PDMPO [2-(4-pyridyl)-5-((4-(2-dimethylaminoethylaminocarbamoyl)methoxy)phenyl)oxazole] or LysoSensor Yellow/Blue DND-160 (Invitrogen L7545) and either 100nM concanamycinA (concA) or an equivalent volume of DMSO for 24 hours (*E. muelleri* n=4 and *S. lacustris* n=3 for each condition). PDMPO fluorescence (ex: 405 nm, em: 490-515 nm) and brightfield images were taken in five 0.408 mm^2^ regions around the periphery of each sponge. The number of spicules per region and the number of spicules per region exhibiting PMDPO fluorescence were counted in ImageJ. The proportion of spicules with PDMPO fluorescence was calculated by dividing number of spicules exhibiting PDMPO fluorescence by the total number of spicules per region. The number of spicules per mm^2^ and the proportion of spicules with PDMPO were averaged across regions to get the final mean values per sponge (raw data provided in Supplementary Table 2, available in our [github repository](https://github.com/maitripaul/Co-option-of-Lysosomal-Machinery-for-Biomineralization/tree/main)). The statistical differences for both aforementioned parameters between sponges treated with concA versus DMSO were calculated using a student t-test. Stereoscope images of all sponges in the experiment were taken before and after treatment to confirm viability of the sponges (Supplementary Figure 6).

## Acknowledgments

We thank Jake Pawley and Haley Womack for their assistance in providing us with sponge gemmules and advice on sponge-rearing. Thank you to the MBL Molecular and Cell Biology of Symbiosis course for kickstarting this collaboration and to its cohort for being unforgettable company. M.R.P. was funded by NSF-GRFP 2023360321.

## Data Availability Statement

Supplementary figures are embedded in main text below. Supplementary tables including supplementary tables 1 and 2 and R code are available in our github repository.

## Supplementary Figures and Table legends

**Supplementary Figure 1.**
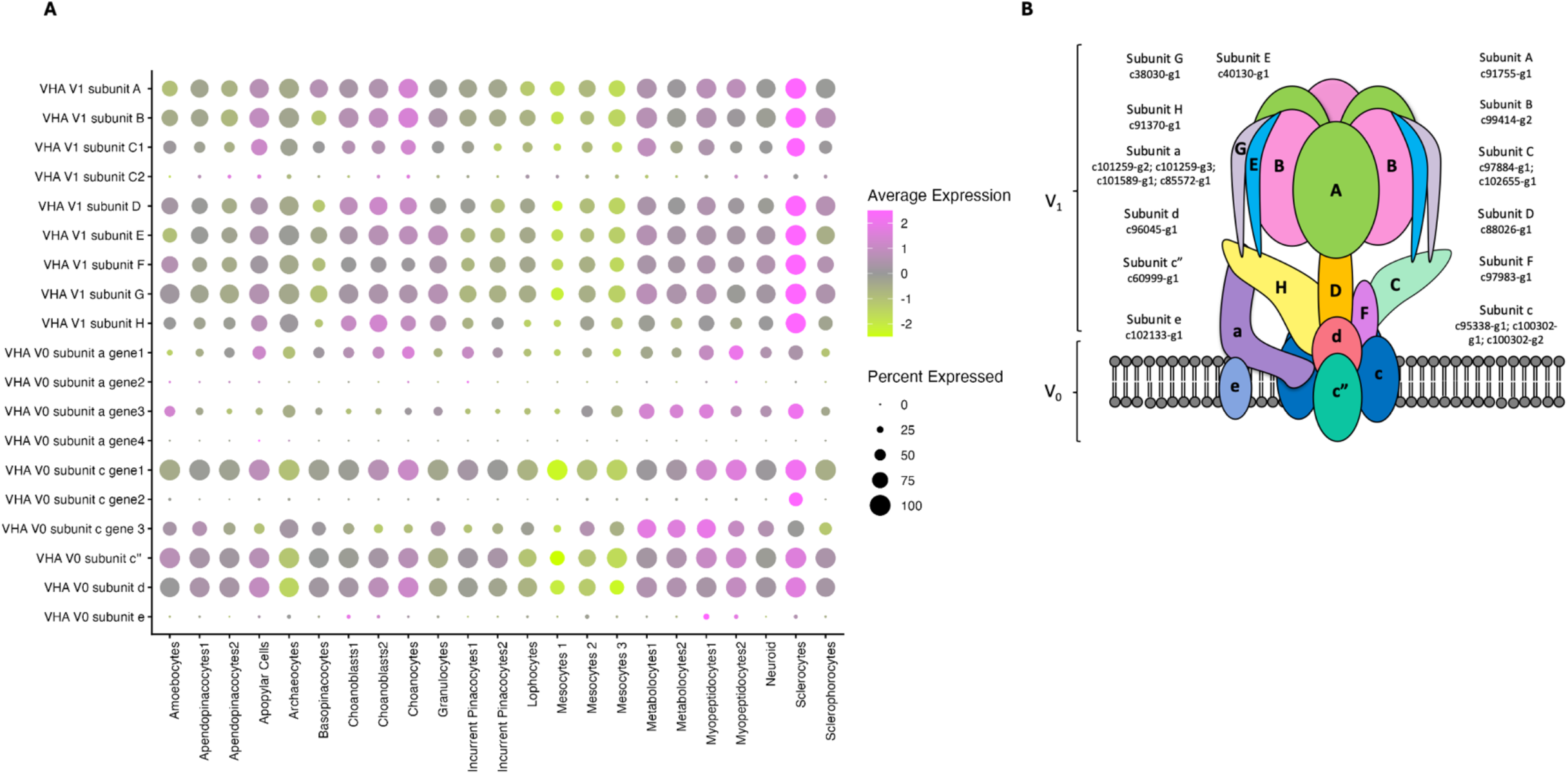
Single-cell RNA-Seq expression of VHA subunits in *S. lacustris*. (A) Heatmap showing expression of genes encoding VHA subunits in *S. lacustris* cell types (scRNASeq data from Musser et al., 2021). Highest expression of almost all VHA subunits occurs in sclerocytes. (B) Cartoon of VHA showing *S. lacustris* genes encoding each subunit.

**Supplementary Figure 2.**
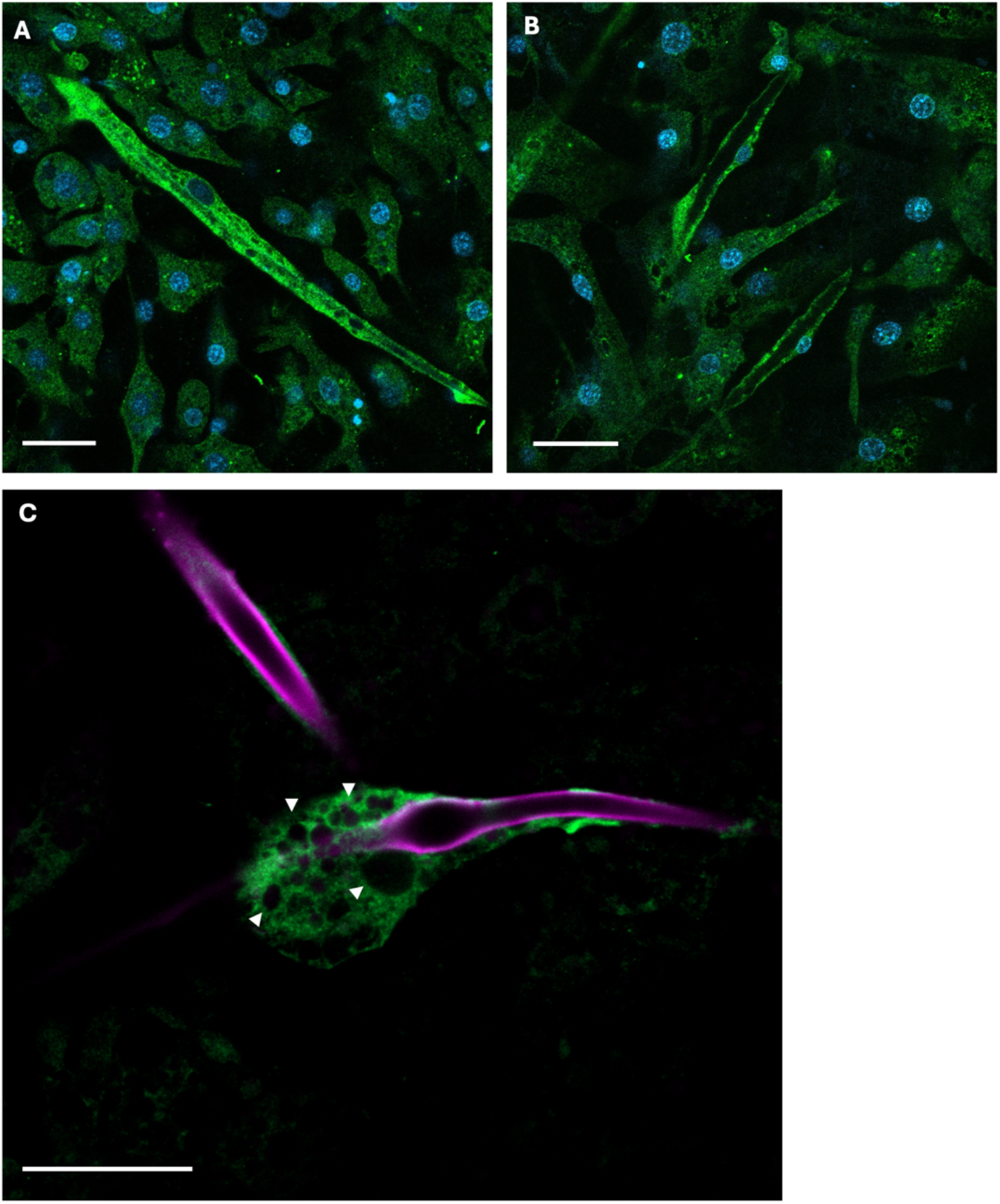
VHA localization at sclerocyte SDV and within cytoplasm in *S. lacustris*. (A, B) Confocal microscopy showing VHA_B_ immunolabeling in *E. muelleri* sclerocytes surrounding spicules and within cytoplasm. (C) Airyscan confocal image showing VHA_B_ localization around *E. muelleri* spicule labelled with PDMPO. Arrows indicate vesicles surrounded by VHA_B_. VHA_B_ in green, DAPI in blue, PDMPO in magenta. Scale bars: 20 μm

**Supplementary Figure 3.**
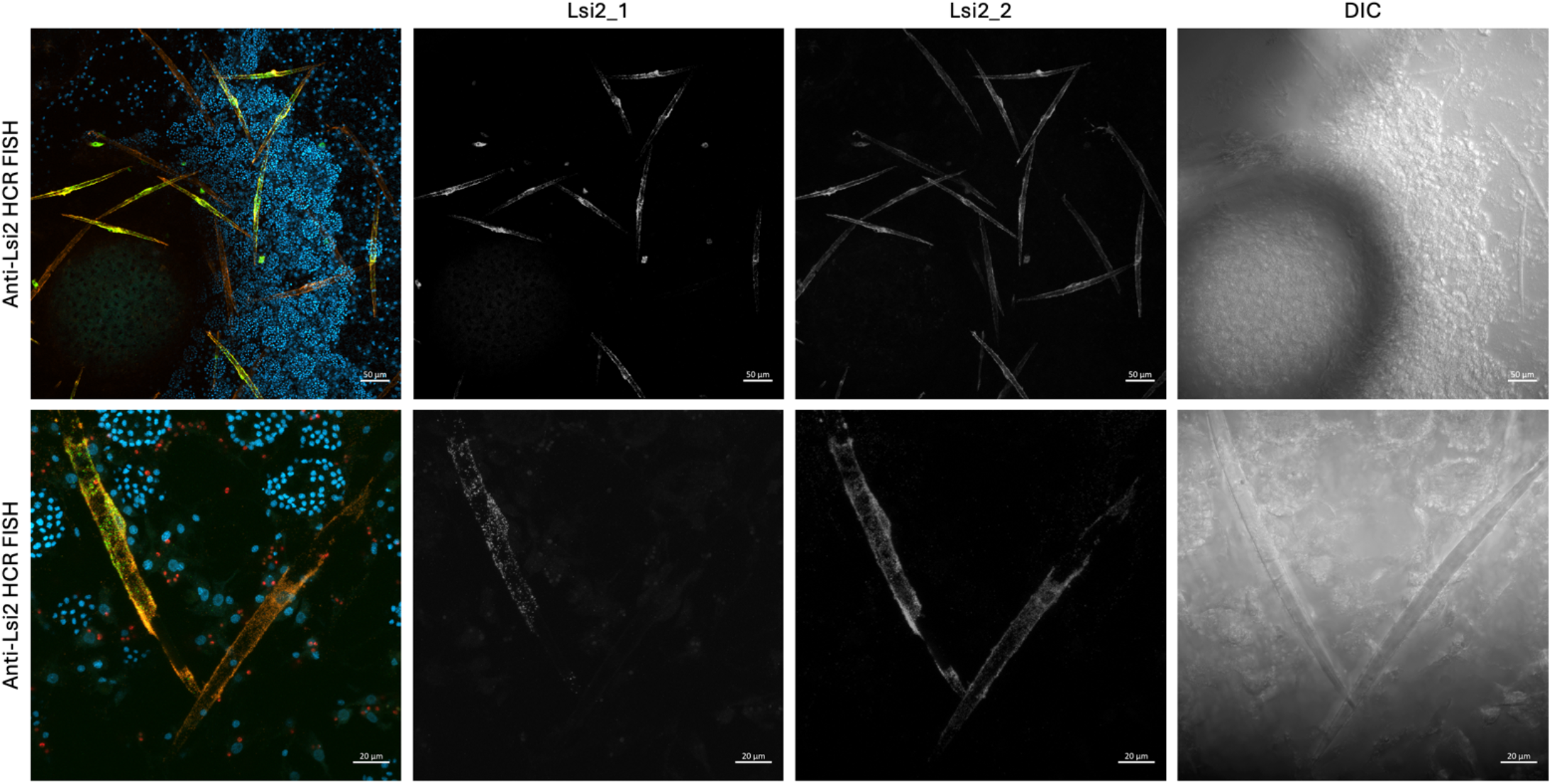
HCR-FISH in *E. muelleri* sponges incubated with probes against Lsi2_1 and Lsi2_2 genes. Representative images showing more ubiquitous Lsi2_2 expression around spicules than Lsi2_1. Lsi2_1 in green, Lsi2_2 in orange, DAPI in blue. Lsi2_1, Lsi2_2, DAPI, and DIC channels are also shown separately in grayscale. Lsi2_1 and Lsi2_2 probes imaged at same laser intensity and gain. All images are orthogonal projections.

**Supplementary Figure 4.**
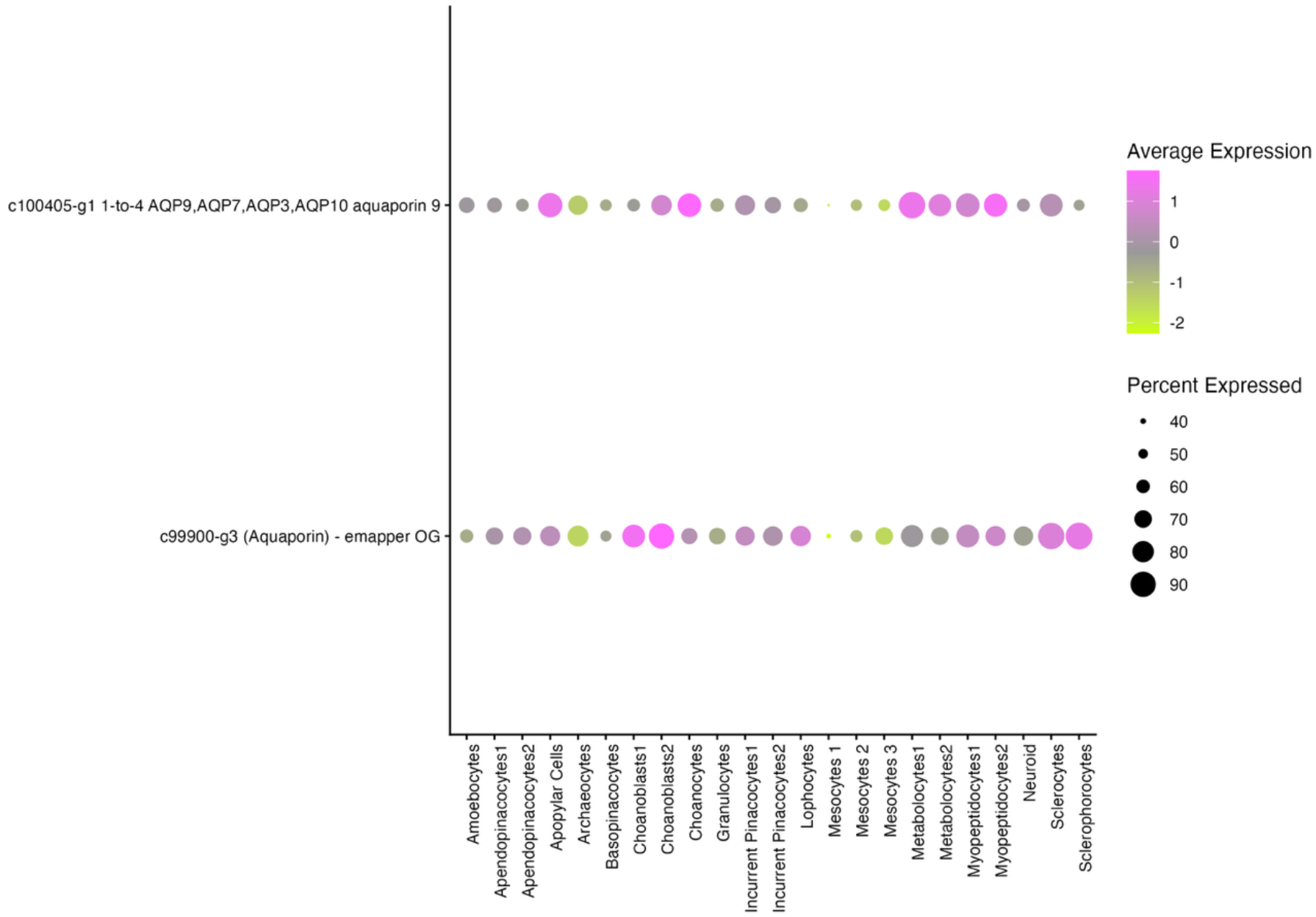
Single-cell RNA-Seq expression of aquaglyceroporins in *S. lacustris*. Heatmap showing expression of genes encoding VHA subunits in *S. lacustris* cell types (scRNASeq data from Musser et al., 2021)

**Supplementary Figure 5.**
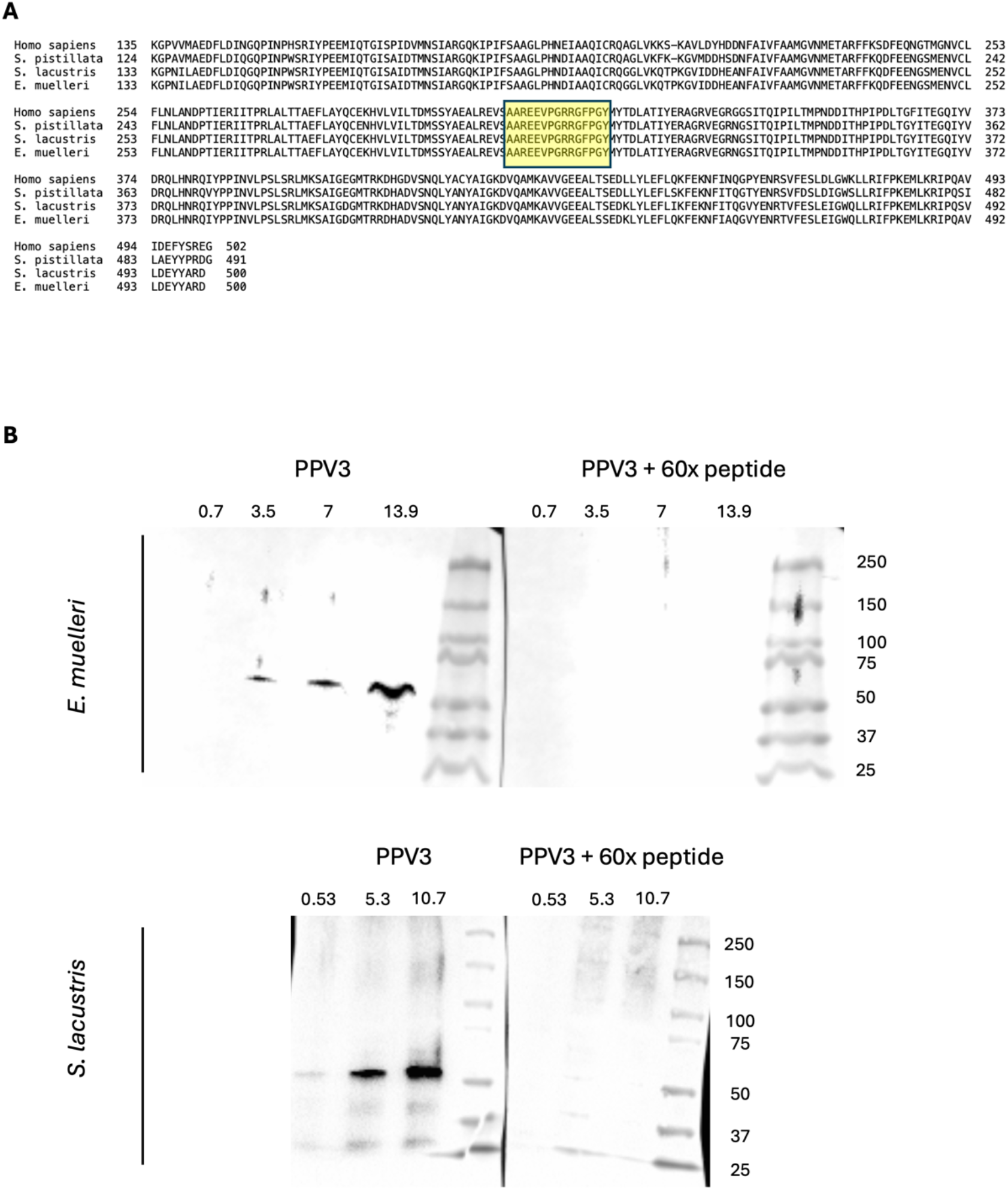
VHA expression and conservation in sponges. (A) Alignment of the human VHA_B_ protein sequence with the coral *Stylophora pistillata, S. lacustris, and E. muelleri* VHA_B_ protein sequences. (B) Western blot detection of VHA in E. muelleri and S. lacustris using anti-VHA_B_ antibody, and anti-VHA_B_ antibody preabsorbed with 60x excess peptide on blots loaded with 0.7 μg, 3.5 μg, 7 μg, and 13.9 μg protein (*E. muelleri*) and 0.53 μg, 5.3 μg, and 10.7 μg protein (*S. lacustris*).

**Supplementary Figure 6.**
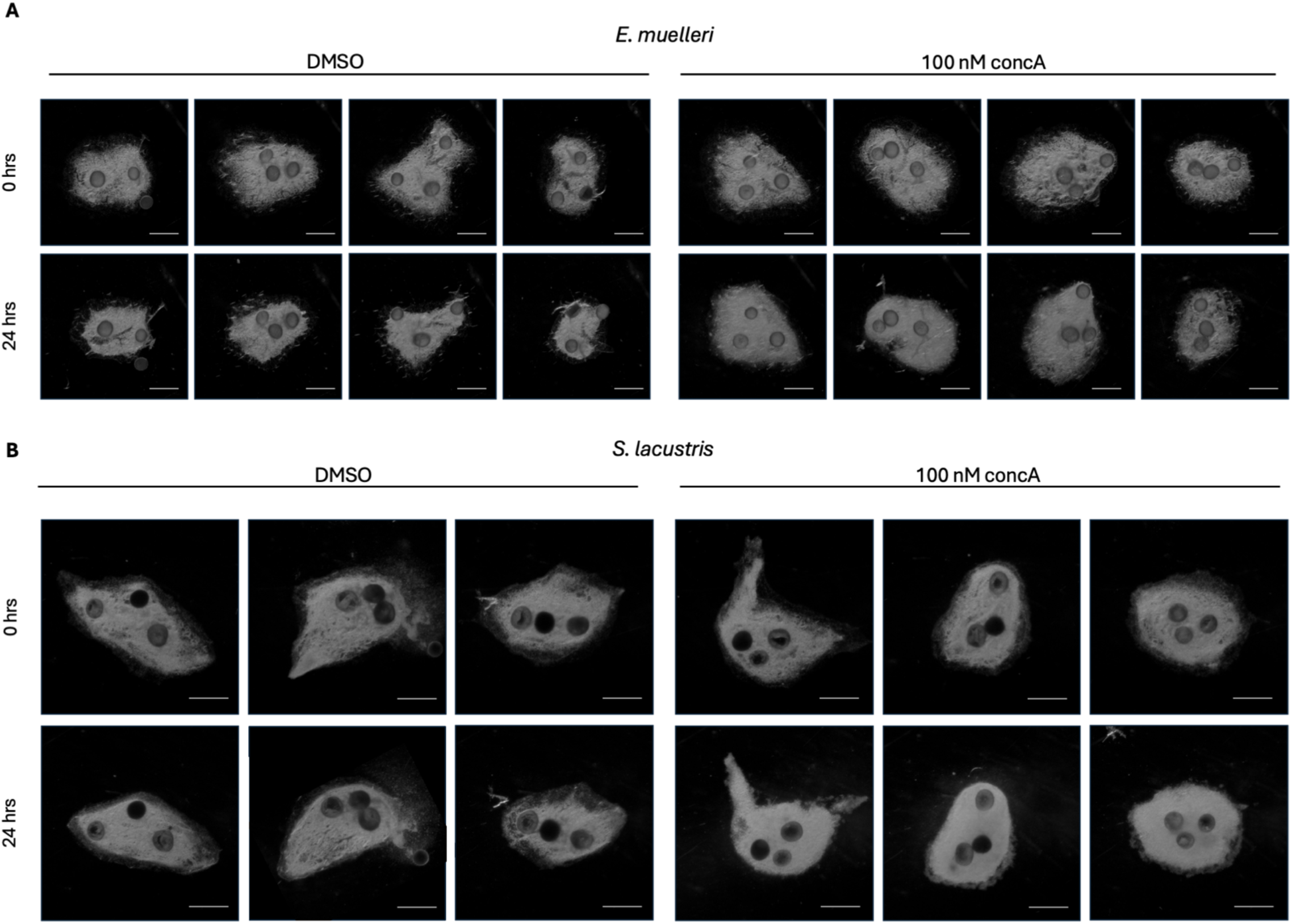
Stereoscope images of *E. muelleri* and *S. lacustris* before and after concA treatment to assess viability. 4-day old sponges before and after 24 hr treatment with 1 uM PDMPO and DMSO or 100 nM concA followed by washout and replacement with spring water + 1uM PDMPO in (A) *E. muelleri* and (B*) S. lacustris*. Scale bars: 1 mm

**Supplementary Table 1** (available at [github repository](https://github.com/maitripaul/Co-option-of-Lysosomal-Machinery-for-Biomineralization/tree/main)): Sequences of *S. lacustris* and *E. muelleri* Lsi2 homologs against which HCR FISH probes were designed

**Supplementary Table 2** (available at [github repository](https://github.com/maitripaul/Co-option-of-Lysosomal-Machinery-for-Biomineralization/tree/main)): Raw data of number of spicules and proportion of spicules with PDMPO fluorescence following concA treatment for Figure 3C, D.

